# Methodological assessment of PDMS passive sampling for skin VOC collection across body sites

**DOI:** 10.64898/2026.07.15.738701

**Authors:** Seibi Kobara, Min Huang, Rizky Ilhamsyah, Daniel Struk, John Dimandja, Peter J. Hesketh, Daniela Chanci Arrubla, Chase Fensore, Carmen Polito, Rishikesan Kamaleswaran, Annette Esper

## Abstract

**Background/Objectives:** Polydimethylsiloxane (PDMS) is a non-invasive and versatile material often used for non-invasive collection of skin-emitted volatile organic compounds (VOCs), with potential applicability in acute and pre-critical care settings. However, most existing PDMS-based methodologies rely on extensive sample preparation and environmental control, limiting their feasibility in time-sensitive clinical contexts.

**Methods:** We conducted a proof-of-concept pilot study in four healthy volunteers to evaluate whether a simplified skin-contact PDMS sampling procedure can capture detectable VOCs and preserve individual-level variation. PDMS strips were applied directly to the skin with minimal preparation, and collected VOCs were analyzed using gas chromatography–mass spectrometry. Donor-associated variability was assessed using Bray–Curtis dissimilarity, and variability in VOC detection was evaluated across body sites.

**Results:** Skin-contact PDMS sampling detected 160 VOCs across four participants. The mean within-donor Bray–Curtis dissimilarity was 0.308, compared with a mean between-donor dissimilarity of 0.347. Preliminary permutation testing showed distinguishable donor profiles (p-value = 0.004). VOC detection variability differed across body sites, with lower coefficients of variation at the forehead, neck, and wrist than at the ankle.

**Conclusions:** Under simplified sampling conditions, skin-contact PDMS captured individual-associated VOC profiles with lower within-donor variability than between-donor variability. These findings support the feasibility of PDMS-based skin VOC sampling in minimally controlled settings. Further validation in larger and clinically relevant cohorts is warranted to assess the utility of PDMS-sampled skin VOCs as potential biomarkers for early disease detection.

## 1. Introduction

Volatile organic compounds (VOCs) are a broad class of organic chemical compounds, characterized by high volatility at room temperature and molecular mobility[1,2]. VOCs are emitted through both natural and anthropogenic processes, including wildfires and industrial activities, and have been associated with immune activation and disease risk [2,3]. Accumulating evidence indicates that VOCs are also continuously emitted by humans through breath and skin, reflecting endogenous metabolism, microbial activity, and environmental exposure [4–6].

Several methodologies have been developed to collect skin-emitted VOCs, including poly-dimethylsiloxane (PDMS), solid-phase microextraction (SPME), cotton pads, Tenax/Carboxen sorbents, and glass beads [6]. These approaches are generally passive and non-invasive, making them well-suited for studying VOC variation across individuals and environments. Nevertheless, substrates such as cotton and porous sorbents may suffer from high environmental signals, batch-to-batch variability, and limited chemical selectivity, which can compromise quantitative reliability[7]. PDMS-based skin-contact sampling offers several advantages, including a high partition coefficient for nonpolar and semi-volatile compounds, low chemical reactivity, and minimal endogenous contamination[8]. The ability to perform solvent-free thermal desorption further simplifies the analytical workflow and reduces analyte loss. In summary, these properties make PDMS an attractive platform for profiling skin-emitted VOCs in both clinical and real-world environments[9].

Critical illnesses such as sepsis and acute respiratory distress syndrome (ARDS) remain leading causes of global morbidity and mortality[10–12], with clinical downstream outcomes strongly influenced by the timeliness of detection and intervention[13]. Despite advances in critical care, early or mild-to-moderate ARDS is frequently underrecognized[11], highlighting the need for methods capable of capturing early physiological dysregulation during pre-critical or rapidly evolving states. In emergency or resuscitation settings, conventional sample collection (e.g., blood draws) may be infeasible or contraindicated, particularly in hemodynamically unstable patients. In this context, skin-contact VOC sampling using PDMS represents a potentially feasible alternative.

However, most existing skin VOC studies rely on extensive PDMS preparation protocols and tightly controlled environmental conditions[6,14–16], which enhance the specificity of VOC capture but limit logistical feasibility in pre-critical and critical settings. In these environments, physiologic changes during resuscitation may provide valuable insight into patient status and prognosis, yet conventional sampling approaches are often impractical. To address this gap, we propose a simplified PDMS-based skin-contact sampling procedure that can be applied and left in place, enabling passive, time-integrated collection of chemical signals during resuscitation. This “set-and-forget” approach captures cumulative exposure rather than a single time-point snapshot. However, the performance of such minimally controlled PDMS-based sampling—particularly with respect to VOC detectability, individual variability, and susceptibility to environmental and material-derived contributions—remains insufficiently characterized.

In this pilot study, we evaluated whether a simplified PDMS-based skin-contact sampling procedure, requiring only minimal skin preparation, can capture VOCs consistent with previously reported human-associated compounds. We further assessed the extent to which clothing, room environment, and material-derived VOCs contribute to the observed signal using parallel control samples. This work is intended as a proof-of-concept to inform future validation and methodological refinement, rather than to generate generalizable biological conclusions.

## 2. Materials and Methods

### 2.1. Study Setting and Participants

We conducted a pilot feasibility study involving skin VOC sampling from four healthy adult volunteers. The study was designed to evaluate the technical feasibility of VOC collection procedures and was descriptive in nature.

This work was not intended to generate generalizable knowledge and case series of five or fewer subjects. This research was reviewed by the Emory Institutional Review Board and determined to qualify as non-human subjects research. All procedures were conducted in accordance with the Declaration of Helsinki.

### 2.2. Skin VOC collection procedure

Polydimethylsiloxane (PDMS) tape (SSP-M823), an FDA-compliant membrane, was purchased from Specialty Silicone Products, Inc (New York, USA)[17]. PDMS strips (3.0mm *×* 15.0mm) were laser-cut using a Trotec Speedy Laser 300 (Trotec Laser GmbH, USA) with Ruby software. Prior to sampling, PDMS strips were thermally conditioned to reduce background volatile contaminants at 356 °F for 60 minutes, allowed to cool to room temperature, and stored in a glass tube, isolated individually.

For proof-of-concept testing, PDMS strips were applied to four body sites: forehead, neck, wrist, and ankle (**Figure 1A**). Before application, participants washed the forehead and hands with soap and paper-dried the skin. PDMS strips were placed directly onto the skin at each site, wrapped by aluminum foil, and secured with 3M Nextcare Bandage (**Figure 1B**). In addition to skin-contact sampling, several PDMS control conditions were prepared, including PDMS attached to clothing on the right thigh (clothing-control), PDMS left exposed in a chemical laboratory environment during the sampling period (room-control), and unexposed blank PDMS (blank-control). Participants then walked outdoors in a university environment for 50 minutes to allow passive absorption of skin-emitted and environmental VOCs.

**Figure 1.**
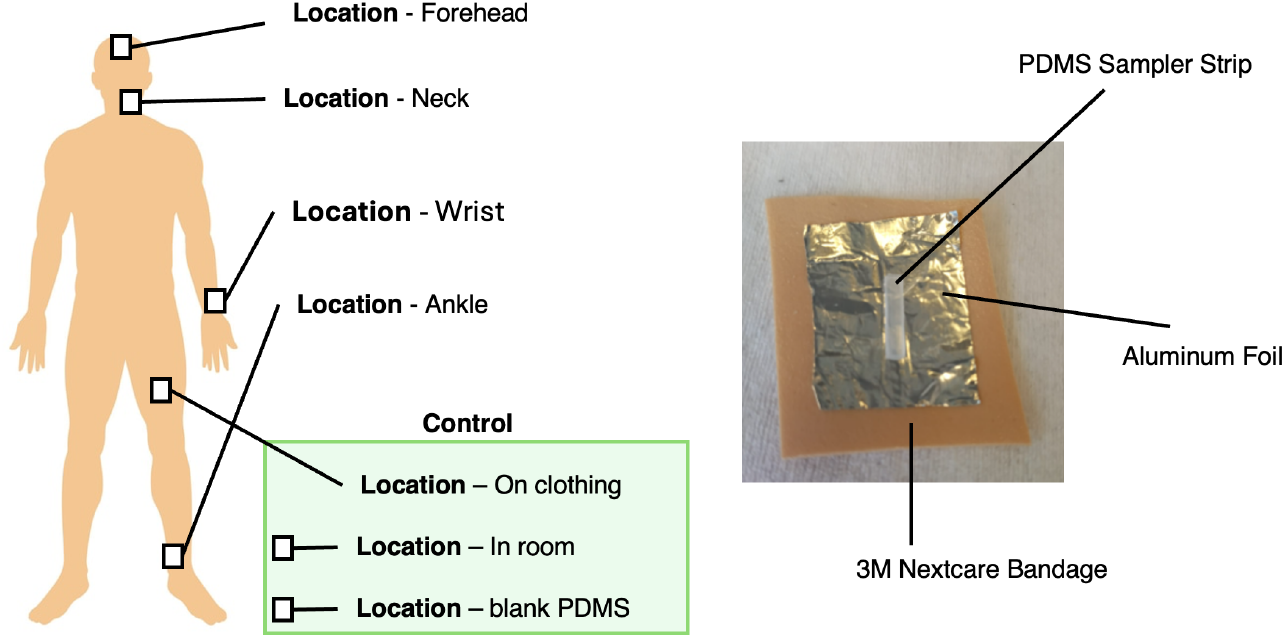
Overview of the experimental design, including the four skin sampling sites, three control conditions, and PDMS tape preparation procedure used for volatile organic compound (VOC) collection.

After sampling, PDMS strips were removed using clean forceps, inserted into a glass inlet, and stored in at -80°C using dry ice until analysis. Collected compounds were analyzed by gas chromatography-mass spectrometry (GC-MS). Detailed GC-MS conditions and quality control procedures are provided in the Supplemental document.

Identified VOCs were mapped to InChIKey identifiers using CIRpy[18], a Python interface to the Chemical Identifier Resolver (CIR) developed by the Computer-Aided Drug Design (CADD) Group at the National Cancer Institute (NCI/NIH). Chemical superclass, class, and subclass annotations were subsequently assigned using the classyfireR[19] package in R.

### 2.3. Statistical analysis

Raw features were detected and aligned across all samples. Features were excluded if they were detected in fewer than four samples. Samples were further filtered if the number of detected compounds was less than 10% of the total number of qualified features retained after feature-level filtering.

To assess inter-individual (donor) and body-site variability, *β* diversity was evaluated using the Bray-Curtis dissimilarity metric. *β* diversity analyses were performed using compound abundance profiles restricted to VOCs detected using skin-contact PDMS. Statistical significance of group differences was assessed using 10,000 permutations.

To evaluate variability in compound detection performance across body sites, the proportion of detected compounds (defined as non-zero peak intensity) was compared among body sites. Statistical significance was evaluated at a two-sided significance level of *α* = 0.05. All preprocessing and statistical analyses were performed using R (version 4.4.1).

## 3. Results

### 3.1. Detected compounds using PDMS

PDMS-based skin VOC sampling was conducted as a proof-of-concept study with healthy adult donors. Four donors were sampled across four body sites (forehead, neck, wrist, and ankle), yielding a total of 13 PDMS samples for analysis; one donor did not undergo forehead and neck sampling for aesthetic reasons, and one sample was removed based on call rate. Following feature detection, alignment, and quality-control filtering, 160 chemical features were retained for downstream analyses. Subsequent analyses focused on evaluating the reliability and reproducibility of PDMS-based skin VOC sampling across donors and body sites.

### 3.2. Donor and body-site variability

Skin-contact PDMS sampling yielded 160 VOCs, including compounds such as acetone, octanal, and decanal (Supplementary Table 1). These detected VOCs were used to assess donor- and body-site–specific variability. Bray–Curtis dissimilarity was computed across all samples and visualized using principal coordinate analysis (Figure 2). The mean within-donor dissimilarity was 0.308, compared with a mean between-donor dissimilarity of 0.347. Permutation testing with 10,000 iterations, in which donor labels were randomly permuted, showing a significant donor-associated effect (p = 0.004).

**Figure 2.**
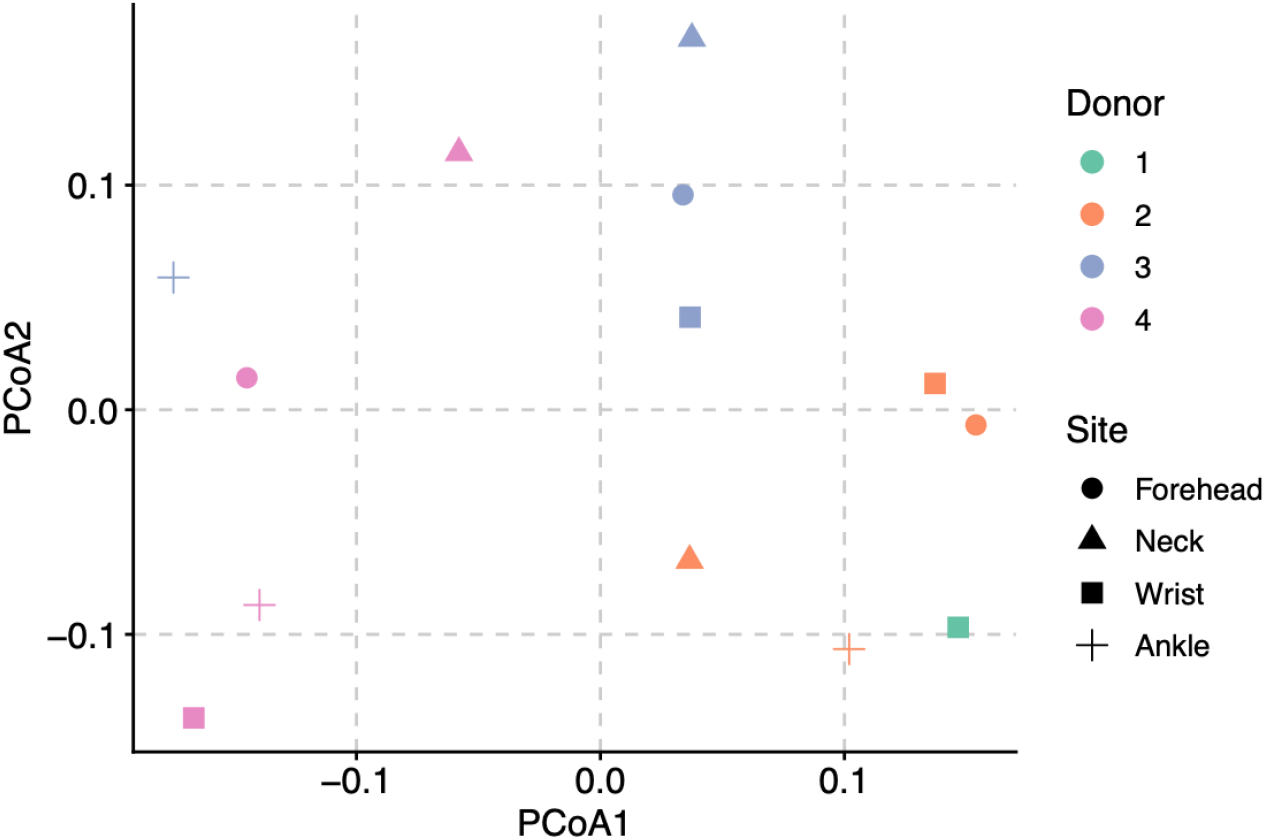
Principal coordinate analysis of Bray-Curtis dissimilarities across donor and body sites for the compound measurements across 160 detected VOCs.

To evaluate the consistency of VOC detection across body sites, the compound abundance matrix was binarized based on the presence or absence of a detectable signal (non-zero peak intensity). The proportion of detected compounds was calculated for each body site. The coefficient of variation (CV) was lowest for the forehead (CV = 0.043), followed by the neck (CV = 0.088), wrist (CV = 0.105), and ankle (CV = 0.615) (Figure 3).

**Figure 3.**
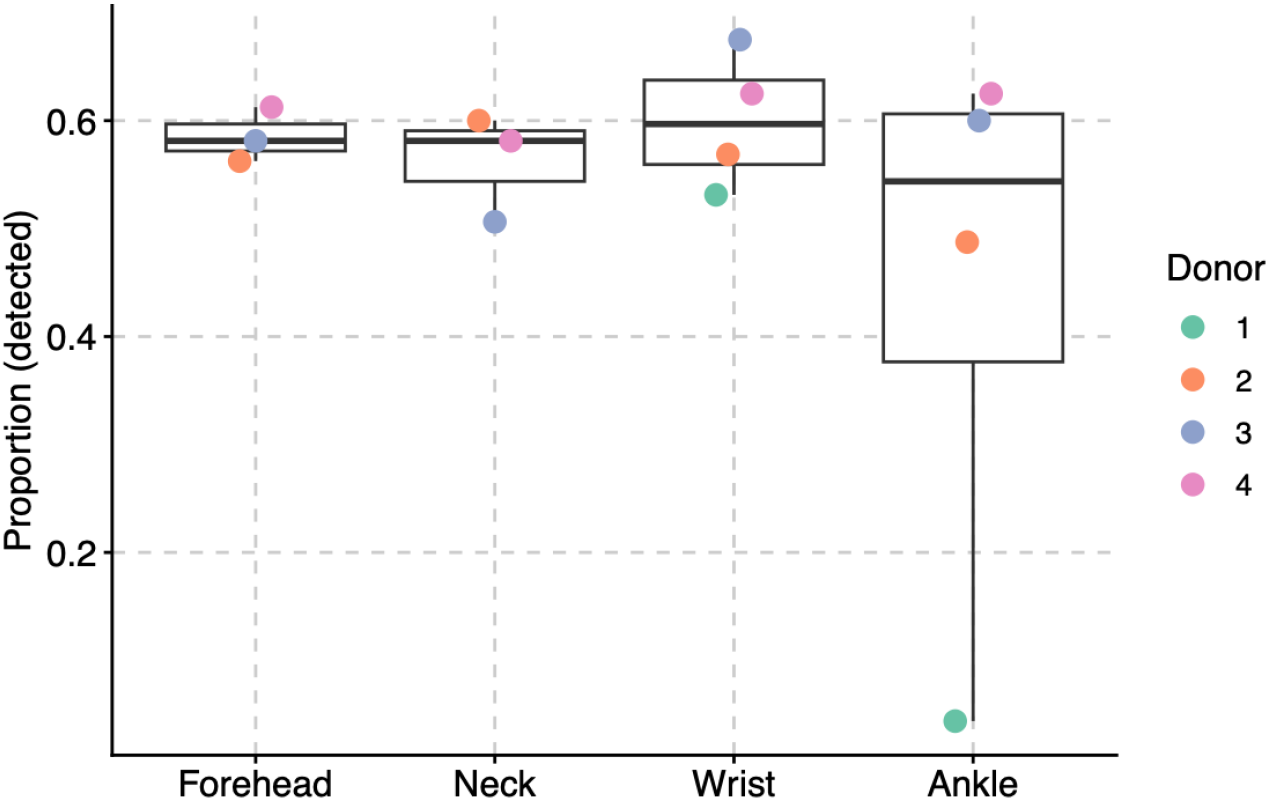
Boxplot showing the proportion of detected VOCs at each skin sampling site (forehead, neck, wrist, and ankle).

### 3.3. Sensitivity analysis using control PDMS samples

We next compared VOCs detected from skin-contact PDMS with VOC profiles obtained from multiple control conditions, including clothing-, room-, and blank-control. In addition, we compared detected VOCs with a curated compilation of previously reported PDMS-derived healthy human-associated VOCs from Mitra et al.[6].

Among VOCs detected using skin-contact PDMS, 94 VOCs were also detected in clothing-control samples, 99 VOCs in room-control samples, and 29 VOCs in blank-control (Figure 4). VOCs overlapping across skin-contact PDMS, clothing-controls, room-controls, and blank-controls included benzothiazoles, aldehydes (e.g., octanal, nonanal, decanal), and fatty acids (e.g., n-decanoic acid). VOCs detected in intersections across experimental conditions are provided in Supplemental Table 1.

**Figure 4.**
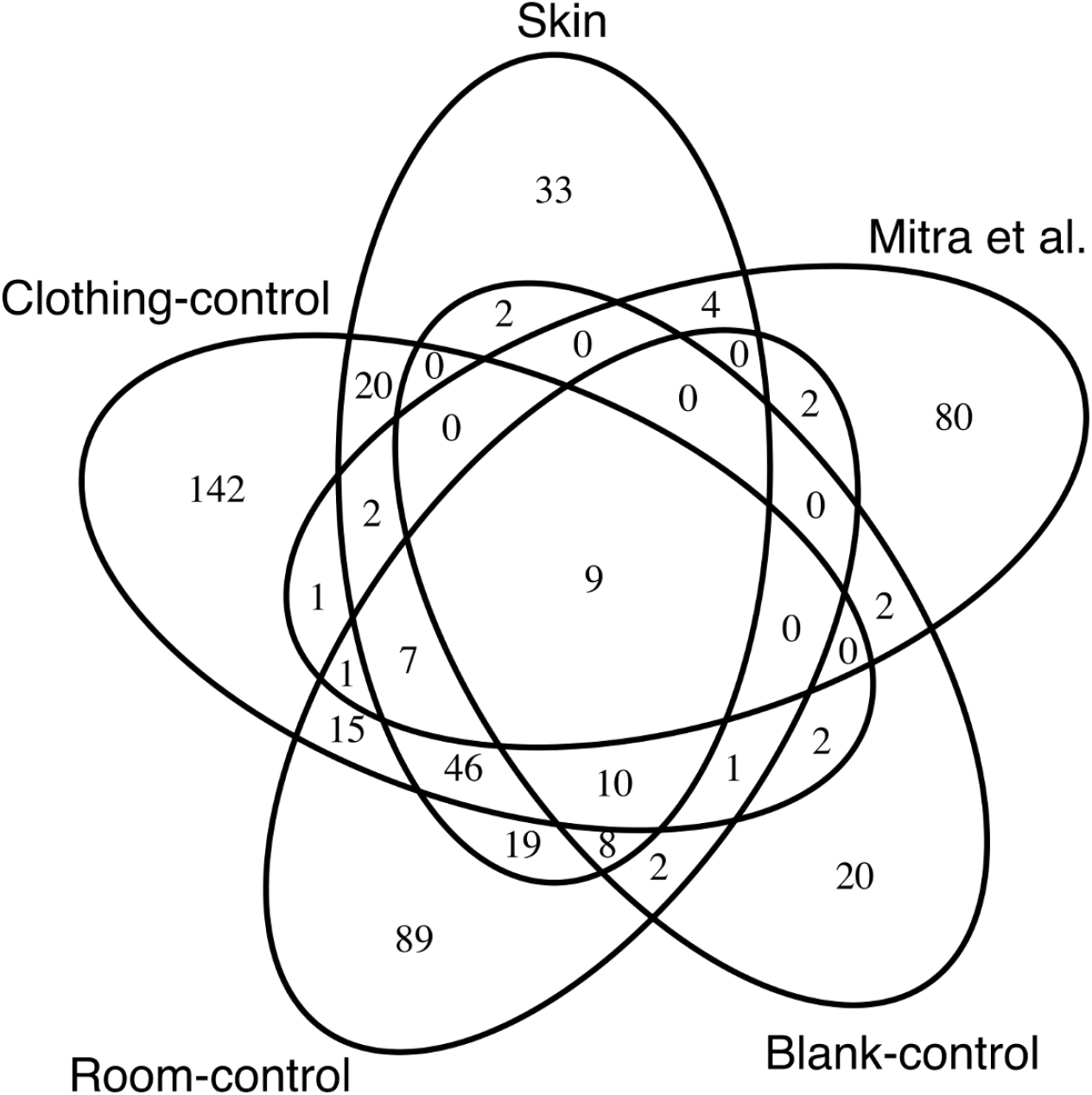
Venn diagram of detected VOCs in PDMS and curated VOCs in literature. Detected represents detected VOCs by skin PDMS, and other control settings, including blank control, room control, and clothing control. The compartment labeled as Mitra et al. represents curated VOCs in literature.

We further compared VOCs detected in skin-contact PDMS with a collection of PDMS-derived healthy human–associated VOCs curated by Mitra et al.[6]. A total of 80 VOCs reported in that compilation were not detected in any experimental condition in our study. These undetected compounds included alcohols and polyols (e.g., dihydromyrcenol and ethanol), short-to medium-chain aldehydes (e.g., heptanal, (E)-2-nonenal, and 2-decenal), short-to medium-chain ketons (2-Nonanone, 2-Decanone, and Cyclopentanone).

## 4. Discussion

In this pilot study, we evaluated the feasibility of using PDMS for the detection of human-associated VOCs under simplified and practical sampling conditions, incorporating multiple quality control procedures. Among the 160 VOCs detected in the primary analysis, within-donor variability was consistently lower than between-donor variability, suggesting that the PDMS-based approach was able to capture reproducible, individual-level VOC profiles. Across body sites, the forehead, neck, and wrist exhibited the lowest coefficients of variation, indicating greater stability of VOC detection at these locations.

PDMS represents a non-invasive, low-burden sampling material with particular relevance for clinical and critical care settings[8,9], where rapid deployment and minimal infrastructure are essential. Although PDMS-based VOC sampling has been widely explored, most prior studies relied on extensive skin-cleaning protocols and tightly controlled environments[6], limiting real-world applicability. In contrast, we evaluated a simplified preparation procedure and explicitly assessed VOC profiles across multiple control conditions. Despite this minimal preparation, the observed donor-specific signal—reflected by significantly lower within-donor than between-donor variability—supports the capacity of PDMS sampling to capture meaningful individual differences in VOC profiles.

Body site selection emerged as an important determinant of reproducibility. While the forehead demonstrated the lowest variability, practical considerations such as cosmetics or topical products may limit its routine use. In contrast, the wrist represented an accessible and potentially more reliable site for PDMS-based VOC sampling in applied or clinical contexts.

Our sensitivity analyses indicate that skin-contact PDMS captures a composite VOC signal influenced by skin-associated chemistry as well as the surrounding microenvironment. Notably, benzothiazoles were detected across experimental conditions, indicating contributions from non-biological sources to skin-associated VOCs[20,21]. While blank controls remain useful for identifying process-related artifacts, the substantial overlap was observed across skin-contact PDMS, clothing-controls, room-controls, and blank-controls, including aldehydes and ketones. Although the direct origin of these compounds cannot be definitively assigned, their presence across multiple conditions suggests complex transfer dynamics, including skin-to-PDMS absorption, ambient air-to-skin deposition, accumulation on clothing, and indirect exchange between these reservoirs[22]. These findings indicate that strict exclusion of overlapping compounds may not be optimal for real-world VOC sampling.

When compared with a curated compilation of PDMS-derived healthy human–associated skin VOCs reported by Mitra et al.[6], 80 of 108 compounds were not detected under any condition in our study. These undetected compounds included alcohols and polyol, medium-chain aldehydes, and short-to medium-chain ketons, which are volatile compounds. In contrast, our skin-contact PDMS sampling uniquely detected compounds such as pentadecanal and 1-hexadecanol, representing a long-chain aldehyde and fatty alcohol, respectively. These compounds are associated with lipid metabolism and oxidative stress[23,24], suggesting that the simplified , passive sampling approach may preferentially capture less volatile, accumulation-prone metabolites at the skin interface.

In future applications targeting acute or pre-critical care settings, individual variability and environmental heterogeneity are expected and largely uncontrollable. Laboratory handling conditions, available sanitizers, personal protective equipment, and emergent medical procedures may vary substantially in urgent clinical scenarios. As such, attempts to completely eliminate exogenous VOC contributions through strict exclusion or subtraction-based filtering are likely impractical. Instead, a more realistic strategy is to develop context-aware machine learning models that integrate real-time environmental monitoring as a dynamic regressor. By treating the environmental background as a high-dimensional feature to be modeled rather than noise to be physically eliminated, we can potentially recover subtle physiological signals from the complex chemical matrix in critical care settings. This study provides the necessary baseline data to train such background-correction algorithms, moving beyond simple subtraction methods toward non-linear source attribution.

Several limitations warrant consideration. This study involved a small number of healthy volunteers and was not designed to evaluate biological or mechanistic inference across individuals. Sampling duration, analytical sensitivity, and compound identification were constrained by the pilot nature of the experiment. Nevertheless, these findings provide a proof-of-concept that informs future study design and highlights the need for validation in larger and clinically relevant cohorts.

## 5. Conclusions

In this pilot study, we quantified VOCs captured by skin-contact PDMS using parallel clothing, room, and blank controls under a skin-cleaning–only preparation. VOC profiles obtained from skin-contact PDMS exhibited significantly lower variability within individuals than between individuals, indicating that this approach can capture reproducible, individual-level VOC patterns. The substantial overlap of VOCs across skin, clothing, room, and blank controls highlights the complexity of VOC transfer and accumulation in real-world settings and suggests that strict exclusion of overlapping compounds may be impractical in applied or acute care contexts. Rather than attempting to eliminate all non-skin contributions, future applications may benefit from computational frameworks that model the dynamic interactions between skin-derived VOCs and environmental or procedural influences across both short- and longer-term exposure timescales.

## Author Contributions

Conceptualization, S.K, M.H., R.I., D.S., P.H., A.E., and R.K.; formal analysis, S.K., M.K., and R.K.; resources, R.I., D.S., J.D., P.H.; writing—original draft preparation, S.K., M.H., and R.K.; writing—review and editing, supervision, R.I., D.S., J.D., P.H., D.A., C.F., C.P., A.E., and R.K.; funding acquisition, A.E. and R.K. All authors have read and agreed to the published version of the manuscript.

## Institutional Review Board Statement

This work was not intended to generate generalizable knowledge and case series of 5 or fewer subjects. This research was reviewed by the Emory Institutional Review Board and determined to qualify as non-human subjects research. All procedures were conducted in accordance with the Declaration of Helsinki.

## Informed Consent Statement

Informed consent was not required due to non-human subjects research.

## Data Availability Statement

R scripts used for data preprocessing and analysis are available from https://github.com/Kamaleswaran-Lab/Skin_VOC_PDMS.git. The datasets underlying this article will be made available from the corresponding author upon reasonable request.

## Acknowledgments

The authors thank Chengkun Yan, Zechary Chou, Vidhi Javia for their early exploration of the work their helpful discussion for this work. S.K. was supported by the National Institutes of Health (NIH) under Award Number R21GM148931 and R01HL170175. R.K. was supported by the NIH under Award Numbers R01GM139967, R21GM151703, R21GM148931, and R01HL170175.

