## Supplementary material for "Methodological assessment of PDMS passive sampling for skin VOC collection across body sites": preprint_sup.pdf

### Supplemental Methods

#### GC-MS Measurement

Gas chromatography-mass spectrometry (GC-MS) analysis was performed using an Agilent 6890 gas chromatography system (Agilent Technologies, Santa Clara, CA) equipped with thermal desorption unit and coupled with a time-of-flight (TOF) mass spectrometer (LECO Pegasus 4D, St. Joseph, MI). Chromatographic separation was achieved on a 30 m Rxi-5 stationary phase column (250 μm ID, 0.25 μm film thickness). Helium (99.999% purity) served as the carrier gas at a constant flow rate of 1 mL/min. Samples were introduced via splitless injection with an initial oven temperature held at 40°C for 30 s, and then increased at a ramp rate of 10°C/min to a final temperature of 250°C and held for another 30 s. The mass spectrometer was operated with an electron ionization energy of 70 eV and an ion source temperature of 230°C. Data were acquired across a mass range of 30–250 amu at an acquisition rate of 200 spectra/sec.

#### Preprocessing

Raw GC–MS data were processed using ChromaTOF software (version 4.72.0.0, LECO Corporation, St. Joseph, MI) with vendor-recommended default settings. ChromaTOF was used for chromatographic peak detection, spectral deconvolution, peak alignment, and compound annotation based on mass spectral matching. Following automated annotation, compounds identified as siloxanes and trimethylsilylated derivatives were removed from downstream analyses because they likely originated from GC column bleed, PDMS-related artifacts, or implausible library assignments.

Compounds detected in fewer than four samples were subsequently excluded, resulting in a final set of 160 compounds. Sample-level call rates were then evaluated, and samples with missing values for more than 10% of the retained compounds were removed. Based on this criterion, one sample corresponding to an ankle skin-contact PDMS collection was excluded from further analyses. Because of the pilot sample size, downstream analyses were based on binary detection status rather than quantitative peak abundance. Therefore, compartment-level VOC summaries reflect whether a putatively annotated compound was detected in at least one sample within each compartment and should not be interpreted as evidence of consistent compound-level enrichment or endogenous origin.
